# Transcriptome-Proteome analysis of human naive and memory B cell subsets reveal isotype and subclass-specific phenotypes

**DOI:** 10.1101/2025.06.05.657894

**Authors:** Jana Koers, Arie J. Hoogendijk, Simon Tol, Floris P.J. Van Alphen, Ninotska I.L. Derksen, Maartje van den Biggelaar, Theo Rispens

## Abstract

Antibodies produced by B cells aid in recognition and clearance of pathogens and is the cornerstone of vaccination strategies. Humans produce nine different antibody isotypes and their effector functions differ according to the type of antigen and route of exposure. Phenotypic variation between isotype-switched B cell subsets is expected but not studied in detail. To obtain a molecular definition of isotype-defined cell identity, we performed proteomics and transcriptomics on isotype-defined populations of human naive and memory B cells (MBCs): CD27^−^IgM^+^IgD^+^, CD27^+^CD38^lo/-^IgM^+^IgD^+^, CD27^+^CD38^lo/-^IgM^+^IgD^−^, and IgA1, IgA2, IgG1, IgG2, IgG3, and IgG4 MBCs (CD27^+^CD38^lo/-^Ig^+^). Combined proteome and transcriptome analysis revealed that mRNA and protein expression profiles separate isotype-defined B cell subsets according to their differentiation status. mRNA and protein expression levels correlated reasonably well for many genes. IgG4-switched B cells were most distinct from naive B cells in terms of mRNA as well as protein expression profiles. Besides a distinct expression profile of cytokine and Fc receptors, we identified a high expression of IgE-coding mRNA in IgG4-switched B cells. SDR16C5 was identified as uniquely upregulated in IgG4-switched B cells. Taken together, this study highlights the distinct phenotypic profile of IgG4-switched B cells.

## Introduction

The ability of B cells to produce antibodies is essential for effective humoral immunity by recognition and clearance of pathogens. During B cell maturation, immunoglobulin (Ig) genes are randomly rearranged to generate B cell receptors (BCRs) which are expressed on the B cell surface. BCRs exist in different isotypes, including IgD, IgM, IgG, IgA and IgE. For IgG, four subclasses are described (IgG1, 2, 3 and 4) and two for IgA (IgA1 and 2). Antigen-naive B cells co-express IgM and IgD BCRs. Upon antigen encounters B cells are able to express other antibody isotypes via a process known as class-switch recombination. For effective humoral responses the expression of the appropriate isotype is essential. The route of antigen exposure and the context in which B cell activation occurs govern selection for a specific isotype which in turn can influence B cell fate and function (1–3).

Activated naive B cells participate in either extrafollicular or germinal center (GC) responses and formation of isotype-switched B cells occurs in both (4–8). Memory B cells (MBCs) can be generated prior to the onset of GCs, or at early and late stages during a GC reaction (9–13). MBCs aid in the generation of immunological memory essential for rapid and effective response to previously encountered pathogens and their related variants (14) and is the cornerstone of vaccination. MBCs are long-lived and recirculate trough the periphery or take up residence in tissues in a quiescent state. Upon secondary antigen exposure, MBCs can undergo further diversification by re-entry into the GC reaction with subsequent isotype-switching, or differentiate into antibody secreting cells in a GC-independent fashion (15–18). B cell fate decision is among others influenced by the expressed isotype and function of MBCs as well as antibody secreting cells is dependent on the expressed isotype (16,17,19,20). Hence, control of isotype expression is relevant to the outcome of both primary and secondary responses.

Naive B cells that predominantly express IgD are anergic and the function of IgD antibodies is still incompletely understood and represent the second least abundant isotype in serum (21,22). IgM is the first antibody to be expressed in the early stages of an immune response in anticipation of affinity-matured antibodies such as IgG and IgA. IgG antibodies are most abundant in serum and confer a central role in systemic immunity and it is often the primary effector antibody raised in response to inflammation, whereas IgA fulfills essential roles in the mucosal immune system. IgE+ B cells are extremely rare in peripheral blood and IgE antibodies mediate allergic reactions and confer powerful effector functions via FcεRs.

Of the four IgG subclasses, IgG4 stands out as it is typically only a minor fraction of an antibody response against bacteria or viruses (55, 56). However, prolonged and/or repeated antigen challenge in the absence of infection can result in a highly IgG4-skewed antibody response, such as during allergen-specific immunotherapy. In this setting, IgG4 titers correlate with relief of allergic symptoms. However, many autoimmune diseases are also characterized by IgG4 autoantibodies. Therefore, possibilities for selective targeting of IgG4-switched B cells may be therapeutically important.

Phenotypic data on B cells expressing different antibody subclasses is sparse and if isotype specificity is also reflected at the cellular level is unclear. Phenotyping studies using flowcytometry previously performed in our lab and by others revealed differences between B cells expressing different IgG subclasses (23–26). IgG4 B cells were found to have reduced expression of chemokine receptor CXCR4 and CXCR5 and complement receptor 2 (CR2/CD21), but higher expression of IgεRII/CD23, compared to IgG1 B cells. Furthermore, IgG4 do not express CCR7, restricting their entry into secondary lymphoid organs and thus most frequently reside within peripheral blood. To obtain a comprehensive and unsupervised molecular definition of isotype and subclass-defined B cell identity, we performed proteomics and transcriptomics for quantitative comparison of human naive and isotype and subclass-defined memory B cells (MBCs).

## Methods

### B cell isolation and cell-sorting

Healthy adult human peripheral blood was obtained with written informed consent in accordance to the guidelines established by the Sanquin Medical Ethical Committee and in line with the Declaration of Helsinki. Donor information is summarized in Supplementary Table 1. Blood samples were drawn outside of the seasonal allergy season. Peripheral blood mononucleated cells (PBMCs) were isolated from fresh buffy coats using Ficoll gradient centrifugation (lymphoprep; Axis-Shield PoC AS). PBMC fractions were screened for adequate yet normal IgG4 MBC frequencies (0.2-1% of total CD19^+^). Total B cells were isolated by untouched magnetic bead separation (Pan B cell isolation kit human; Miltenyi Biotec) and studied subsets were separated using flow cytometric cell-sorting based on expression of CD19, CD27, CD38 and BCR isotype (FACS Aria III and FACS Aria IIIu, BD Biosciences). During sorting cells that expressed multiple BCRs isotypes, due to promiscuous antibody staining or recent class switch events were excluded.

### Antibodies

The following antibodies were included: anti-CD19 (clone SJ25C1, 563325), anti-IgD (clone IA6-2, 561315), and anti-IgM (clone G20-127, 562618) from BD Biosciences. Anti-IgA1 (clone B3506B4, 9130-30), anti-IgA2 (clone A9604D2, 9140-31), and anti-IgG2 (clone HP6002, 9070-02) from SouthernBiotech. Anti-IgG1 (clone HP6188, M1325), Anti-IgG3 (clone HP6095, M1270), and anti-IgG4 (clone HP6098, M1271) from Sanquin Reagents. Anti-CD27 (clone O323, 63-0279-42), anti-CD38 (clone HB7, 25-0388-42), and Live/Dead Fixable Near-IR Dead cell stain (L10119) from ThermoFisher scientific. Each antibody was titrated to optimal staining concentration using PBMCs.

### Sample preparation for mass spectrometry

FACS-separated B cell subsets from 8 donors were sorted into Protein LoBind tubes (Eppendorf), and washed with PBS. For IgG4 four donors were pooled to reach sufficient cell numbers for proteomics analysis (n = 4) and for IgG3 two donors were pooled (n = 6) (Supplementary Table 2). Tryptic peptides were prepared similarly to a previously described method (28). Briefly, depending on the acquired cell number per sample (Supplementary Table 2) cells were lysed in 10-25uL 1% Sodium Deoxy Cholate (SDC) (Sigma Aldrich, Germany) 10mM TCEP (Thermo Scientific, USA), 40mM ChloroAcetamide (Sigma Aldrich, Germany), 100mM TRIS-HCl pH 8.0 (Life Technologies, UK), boiled at 95 °C for 5 minutes and sonicated for 10 minutes in a BioRuptor Pico (Diagenode, Belgium). A double volume of 100 mM TRIS-HCl pH 8.0 was added, containing 125-325ng Trypsin/LysC (Thermo, USA). Samples were digested overnight at 25°C. Next, samples were acidified by addition of 1% (v/v) trifluoroacetic acid (Thermo Scientific, USA), centrifuged and supernatants containing the peptides were loaded on in-house prepared SDB-RPS STAGEtips (Empore, USA). STAGEtips were washed with 0.1% TFA and peptides were eluted in 5% (v/v) ammonium hydroxide (Sigma Aldrich, Germany), 80% v/v acetonitrile (BioSolve). Sample volume was reduced by SpeedVac and supplemented with 2% acetonitrile, 0.1% TFA to a final volume of 10 μL. 3 μL of each sample was injected for MS analysis.

### Mass spectrometry data acquisition

Tryptic peptides were separated by nanoscale C18 reverse phase chromatography coupled on line to an Orbitrap Fusion Tribrid mass spectrometer (Thermo Scientific) via a nanoelectrospray ion source (Nanospray Flex Ion Source, Thermo Scientific). Peptides were loaded on a 20 cm 75–360 µm inner-outer diameter fused silica emitter (New Objective) packed in-house with ReproSil-Pur C18-AQ, 1.9 μm resin (Dr Maisch GmbH). The column was installed on a Dionex Ultimate3000 RSLC nanoSystem (Thermo Scientific) using a MicroTee union formatted for 360 μm outer diameter columns (IDEX) and a liquid junction. The spray voltage was set to 2.15 kV. Buffer A was composed of 0.1 % formic acid and buffer B of 0.1 % formic acid, 80% acetonitrile. Peptides were loaded for 17 min at 300 nL/min at 5% buffer B, equilibrated for 5 minutes at 5% buffer B (17-22 min) and eluted by increasing buffer B from 5-15% (22-87 min) and 15-38% (87-147 min), followed by a 10 minute wash to 90 % and a 5 min regeneration to 5%. Survey scans of peptide precursors from 400 to 1500 m/z were performed at 120K resolution (at 200 m/z) with a 4 × 105 ion count target. Tandem mass spectrometry was performed by isolation with the quadrupole with isolation window 1.6, HCD fragmentation with normalized collision energy of 30, and rapid scan mass spectrometry analysis in the ion trap. The MS2 ion count target was set to 1.5 × 104 and the max injection time was 35 ms. Only those precursors with charge state 2–7 were sampled for MS2. The dynamic exclusion duration was set to 60 s with a 10 ppm tolerance around the selected precursor and its isotopes. Monoisotopic precursor selection was turned on. The instrument was run in top speed mode with 3 s cycles.

### RNA Extraction

FACS-separated B cell subsets from 8 donors were sorted into DNA LoBind tubes (Eppendorf) and washed with PBS. Total RNA was isolated from B cell sorted subsets using Qiagen RNeasy Plus Universal mini kit according to the manufacturer’s instructions (Qiagen, Hilden, Germany). RNA samples were quantified using Qubit 4.0 Fluorometer (Life Technologies, Carlsbad, CA, USA) and RNA integrity was checked with RNA Kit on Agilent 5300 Fragment Analyzer (Agilent Technologies, Palo Alto, CA, USA). Total RNA samples that had inadequate quantity or an RNA Integrity Number (RIN) < 8 were excluded from the library preparation.

### RNA Library Preparation and Sequencing

RNA sequencing library preparation was prepared using NEBNext Ultra II Directional RNA Library Prep Kit for Illumina following manufacturer’s instructions (NEB, Ipswich, MA, USA). Briefly, mRNAs were first enriched with Oligo(dT) beads. Enriched mRNAs were fragmented. First strand and second strand cDNA were subsequently synthesized. The second strand of cDNA was marked by incorporating dUTP during the synthesis. cDNA fragments were adenylated at 3’ends, and indexed adapter was ligated to cDNA fragments. Limited cycle PCR was used for library amplification. The dUTP incorporated into the cDNA of the second strand enabled its specific degradation to maintain strand specificity. Sequencing libraries were validated using NGS Kit on the Agilent 5300 Fragment Analyzer (Agilent Technologies, Palo Alto, CA, USA), and quantified by using Qubit 4.0 Fluorometer (Invitrogen, Carlsbad, CA). The sequencing libraries were multiplexed and loaded on the flowcell on the Illumina NovaSeq 6000 instrument according to the manufacturer’s instructions. The samples were sequenced using a 2×150 Pair-End (PE) configuration v1.5. Image analysis and base calling were conducted by the NovaSeq Control Software v1.7 on the NovaSeq instrument. Raw sequence data (.bcl files) generated from Illumina NovaSeq was converted into fastq files and de-multiplexed using Illumina bcl2fastq program version 2.20. One mismatch was allowed for index sequence identification.

### Data processing and Analysis

Raw MS data files were acquired with XCalibur software (Thermo Fisher Scientific) and processed with MaxQuant 1.6.2.10 software (29). Peptides were searched against the homo sapiens Uniprot database (downloaded March 2019, 73,932 entries). Enzyme specificity was set to trypsin with a maximum of 2 missed cleavages, and carbamidomethylation on cysteine residues was used as fixed modification. The false discovery rate (FDR) was 0.01 for peptide and protein levels with a minimum length of 7 amino acids. Label-free quantitation (LFQ) was performed with a minimum ratio count of 2 based on unique peptides for quantification. MaxQuant output files were loaded in R 4.1.2 for further analysis. Proteins were filtered by ‘potential contaminant’, ‘reverse’ and ‘only identified by side’ and LFQ intensities were log2 transformed. Statistical significance was determined with moderated t-tests using the limma package (30). A Benjamini-Hochberg multiple testing corrected P-value <0.05 and absolute log_2_ fold change of >1 was considered statistically significant and relevant. RNA fastq sequencer output quality was determined using FastQC and transcripts were aligned to the human hg38 genomic reference (release 104) sequence using STAR (31). Differential expression analysis was performed using DESeq2 (32), applying a significance threshold of a Benjamini-Hochberg multiple testing corrected P*-*value of < 0.05 and absolute log2 fold change of >1. Correlation between data and disorder specific theoretical protein profiles was performed using Pearson correlation. Transcriptomes and proteomes were merged based on ENSEMBL identifiers and co-expression analysis was performed using the WGCNA package (33). Gene ontology and pathway overrepresentation analyses were performed using the clusterprofiler package (34).

### Statistical analysis

Differences between groups were analyzed using a one-way ANOVA and Tukey’s multiple comparison test (each group against every other group). A p value <0.05 was considered significant. Statistical test, excluding those evaluating RNAseq and proteomics data, were performed using GraphPad Prism 9.1.1.

### Data availability

The mass spectrometry data have been deposited to the ProteomeXchange Consortium (http://proteomecentral.proteomexchange.org) via the PRIDE partner repository with dataset identifier PXD060953. RNAseq data is deposited to the GEO repository and can be accessed with accession number GSE298777.

## Results

### Transcriptomics and proteomics of human peripheral blood naive and subclass-defined memory B cells

To define subclass-defined MBC identity at the molecular level we performed quantitative proteomics and transcriptomics (Fig. 1). To this end, nine CD19^+^ B cell subsets were isolated from peripheral blood mononuclear cells (PBMCs) of healthy donors (n = 16). These subsets comprised naive B cells (CD27^−^IgM^+^IgD^+^), non-switched MBCs (CD27^+^ CD38^lo/-^IgM^+^IgD^+^), IgM MBCs (CD27^+^ CD38^lo/-^IgM^+^IgD^−^), and isotype switched IgA1, IgA2, IgG1, IgG2, IgG3, and IgG4 MBCs (CD27^+^ CD38^lo/-^ IgA1-2^+^/IgG1-4^+^; Fig. 1A). CD27^+^ B cells that express IgE BCRs are extremely rare in human peripheral blood (35) and were therefore not included in this study. Due to the low frequency of IgG4 B cells we did not further segregate memory B cell subsets using additional phenotypic markers (36). RNA sequencing on the nine B cell subsets (n = 8) identified in total 60,664 unique mRNAs. High-resolution mass spectrometry (MS, identified 3511 proteins across all subsets (n = 8, Fig. S1B). For each subset the highest BCR isotype expressed was in accordance to the isolation strategy validating our approach (Fig. S1C).

**Figure 1.**
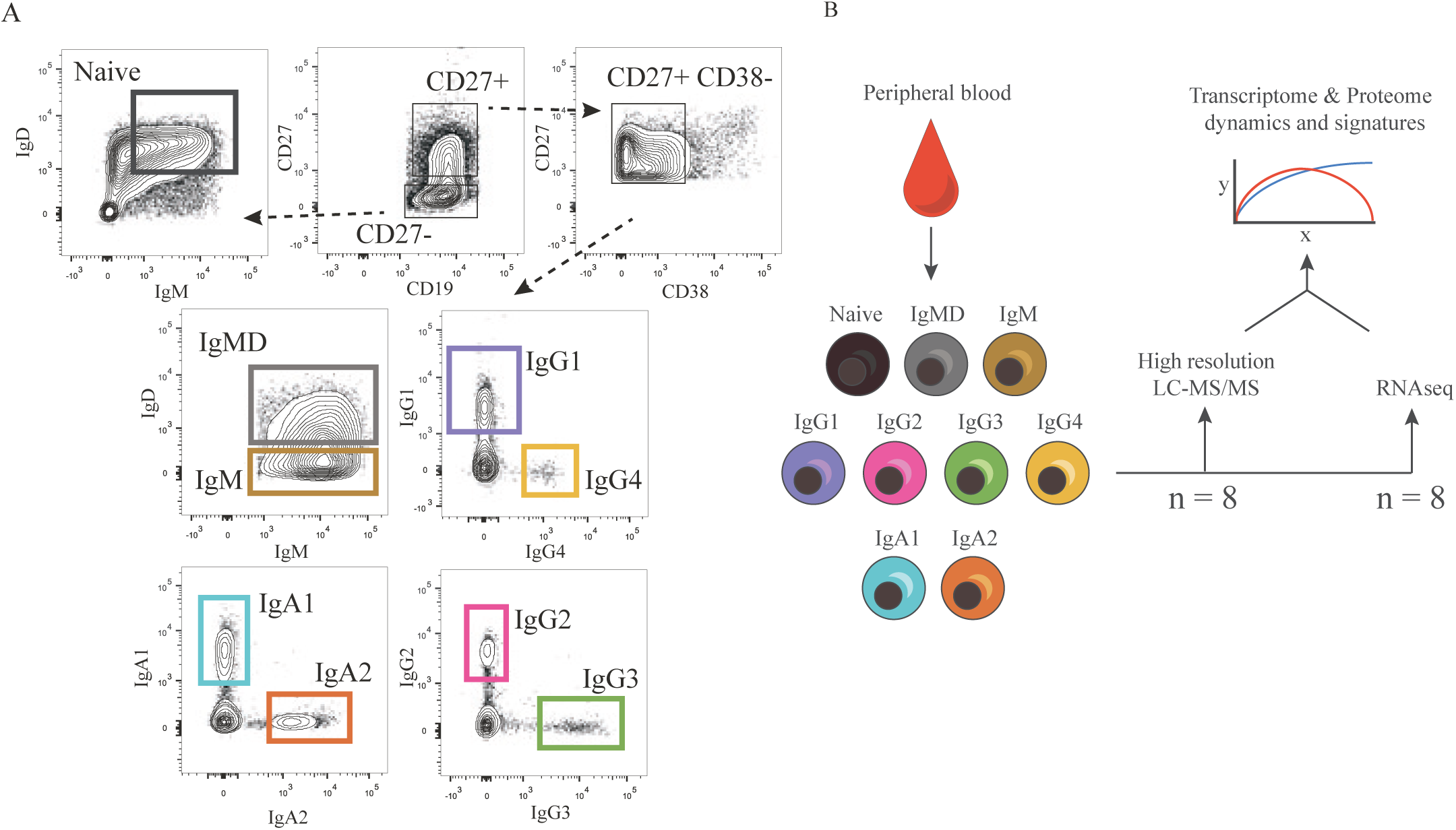
Transcriptomics and Proteomics of human peripheral blood naive B cells and Ig-isotype defined memory B cells. (**A**) Representative flow cytometric profiles of isolated human B cells subsets. (**B**) Schematic representation of experimental set-up and analysis. Naive B cells and memory B cell (MBC) subsets were isolated by gradient centrifugation and flow cytometry and analyzed by liquid chromatography-tandem mass spectrometry (MS) and RNA sequencing.

### Gradual B cell differentiation according to isotype identity

Principal component analysis demonstrated a highly similar pattern in relatedness between B cell subsets for transcriptomics and proteomics. Naive B cells separated from IgMD^+^ and IgM^+^ MBCs, and were distinct from switched MBCs (Fig. 2A). The different types of switched memory B cell subsets showed limited separation in PC1, suggesting phenotypic resemblance of these subsets. The order of B cell subsets along PC1 resembled the order of isotypes found on the Ig locus, whereas PC2 confirmed the close relatedness of biological replicates (Fig. S2B). IgG4 MBCs were most distinct from naive B cells and clustered the opposite end of PC1 and also exhibited the highest number of significantly different mRNAs and proteins (Fig. 2B, S2A).

**Figure 2.**
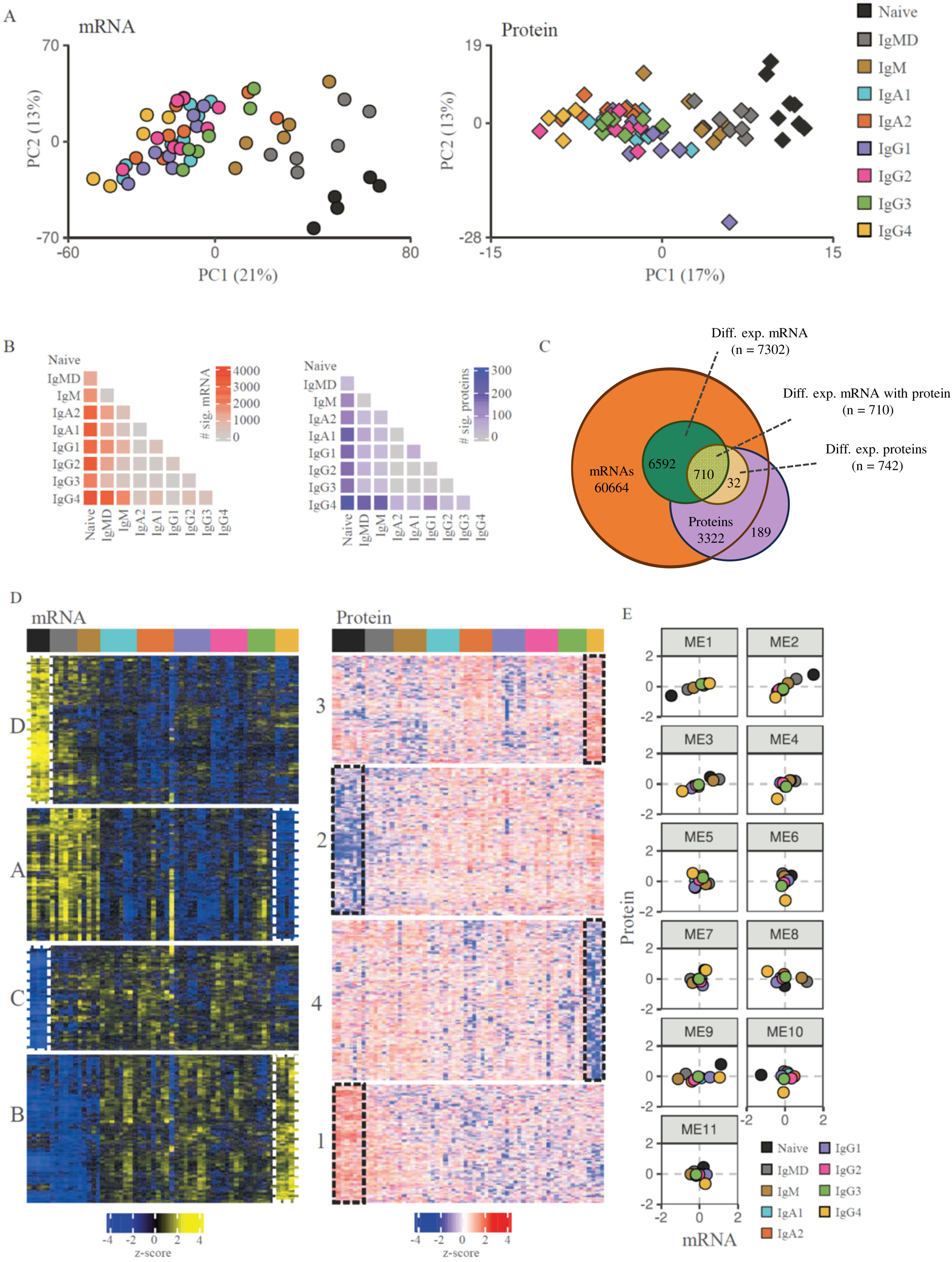
Transcriptomics and proteomics profiles of B cell subsets. (**A**) PCA plot of transcriptomes (left) and proteomes (right). Each color represents a subset and each symbol a single donor. (**B**) Heatmaps display the number of significantly differently expressed mRNAs (left) or proteins (right). (**C**) Venn diagram showing expressed mRNA, differentially expressed mRNA, quantified proteins, and differentially expressed proteins. Note that for most proteins we detected mRNAs, but not vice versa. (**D**) K-means clustering showing relative mRNA (left) and protein (right) expression levels (z-score) of the differentially expressed genes (n = 7308, p < 0.05) and proteins (n = 738, FDR < 0.05) among B cell subsets. Colored columns respond to different subsets (n = 4-8). Letters (mRNA) and numbers (protein) indicate clusters. Dashed white (mRNA) or black (protein) lines represent distinct expression signatures for naive B cells and IgG4 MBCs. (**E**) Co-expression modules depicting mRNA and protein expression levels for all transcript-protein pairs.

K-means clustering of the differentially expressed mRNAs (#7302) and abundant proteins (#742) subdivided both datasets in 4 clusters with distinct expression profiles (Fig. 2C-D). mRNA clusters C & D and protein clusters 1 & 2 are driven by a high or low expression of genes in naive B cells. Proteins upregulated in naive B cells (cluster 2) included B cell identity transcription factors (PAX5, BACH2, and IRF8) and BCR signaling proteins (CD79A/B, CD22, BLNK). Gene Ontology (GO) enrichment analysis indicated cluster D to comprise pathways involved in B cell activation (Suppl. Table 3, 4). Clusters A & B and 3 & 4 were driven by high and low expression of genes in IgG4 MBCs, further discussed below.

For most identified proteins the corresponding mRNA was also identified (#3322 = 95%, Fig. 2C). Of these, 955 were significantly differentially expressed between the nine B cells subsets on either mRNA or protein level. A weighted correlation network analysis was performed to elucidate transcript-protein dynamics and identify modules (ME) of protein and RNA with correlating expression patterns (Fig. 2E) (33,37). This resulted in 11 modules (MEs), ranging in size from 28 to 479 transcript-proteins pairs.

In the module containing the largest fractions (68%) of transcript-protein pairs (ME1, 2 and 3), mRNA and protein dynamics were in concordance. The remaining (32%) of transcript-protein pairs showed varying discordance between mRNA and protein dynamics (ME4-ME11). Most prominently, in ME8 and ME9 RNA expression dropped or increased while protein abundance remained similar. This may reflect proteins with a long half-life or protein storage.

GO enrichment analysis was used to identify molecular functions describing the transcript-protein pair MEs. ME2 describes transcript-protein pairs involved in B cell activation, proliferation, and BCR signaling and were highest expressed in naive B cells and lowest in IgG4 MBCs. For ME5 GO enrichment revealed as molecular function “MHC class II activity” and indeed contained transcript-protein pairs for genes concerning antigen presentation and the formation of immunological synapses, like CD79B, LAT2 and several HLA class II proteins. In summary, comprehensive proteome and transcriptome analysis of naive and MBC subsets in human peripheral blood revealed that mRNA and protein expression profiles separate B cell subsets according to differentiation status, correlated for many genes, and IgG4 MBCs were most distinct from naive B cells.

### Common signatures for naive B cells versus MBCs

Prompted by the observation of clusters predominantly driven by up or downregulation of mRNAs and proteins in naive B cells, we performed a supervised classification to identify mRNAs and proteins that closely follow this dynamics of a gradual up or downregulation of mRNAs and proteins from naive B cells via IgMD MBCs, IgM MBCs and switched MBCs, similar to what was observed for PC1 in the PCA. (Fig. 3A). Supplementary Table 5 and 6 show for each B cell subset upregulated mRNAs and proteins (>0.6) and downregulated mRNAs and proteins (<-0.6). For all subsets we identified corresponding IGH isotype among upregulated mRNAs and proteins, which validates our approach. Note that we could not identify unique peptides for IGHG1, IGHG2 and IGHG3 due to high homology, therefore no intensity value could be assigned to these proteins. For most subsets we identified several specifically up- or downregulated mRNAs, but fewer proteins.

**Figure 3.**
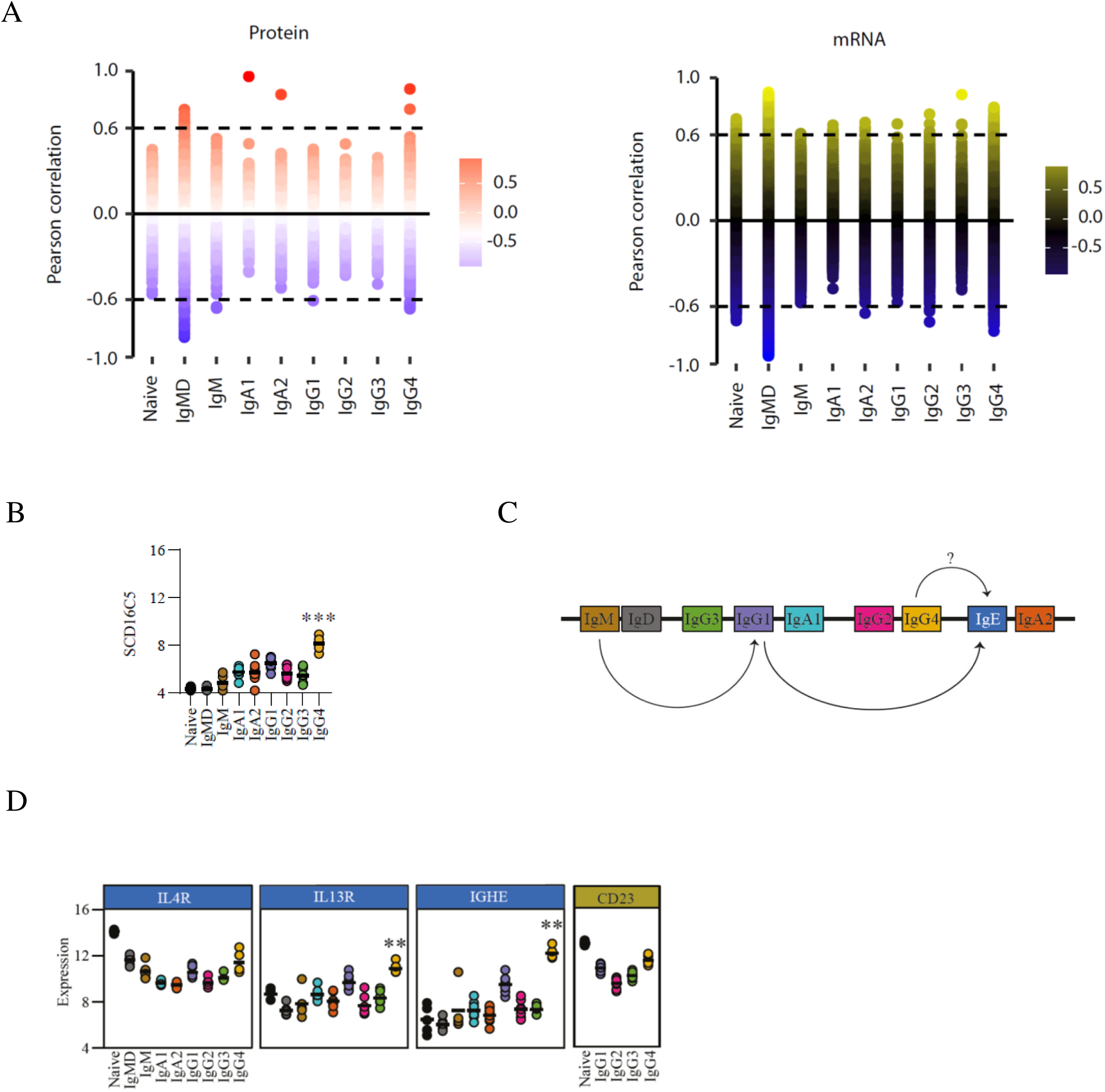
Protein/mRNA signature of IgG4-swiched B cells. (**A**) B cell subset-specific up or downregulated mRNAs and proteins. Perfect fit analysis of mRNAs or proteins for each B cell subset (p <0.05). Dashed lines depict cut-off for upregulation >0.6 and cut-off for downregulation <0.6 of mRNAs and proteins for a subset compared to all other B cell subsets. (**B**) mRNA expression levels of SDR16C5. (**C**) Schematic representation of the human IGH constant gene regions. Arrows indicate (potential [?]) sequential isotype switching events. (**D**) mRNA expression levels of IL4R, IL21R, IGHE and CD23.

The common naive B cell protein signature consisted of 17 proteins with higher levels, and 41 proteins with lower levels in naive B cells compared to MBCs (Supplementary Table 6). This signature included surface receptors IGHD, CD22 and CD72, BCR-associated lipid raft protein RFTN1, and AKT mediator TCL1A, all expressed in high abundance compared to MBCs, as well as signaling regulators Galectin-1, DOK3 and THEMIS2, integrins ITGB1 and ITGAM, and myeloid cell nuclear differentiation antigen MNDA, all expressed at low abundance in naive B cells compared to MBCs. This naive B cell signature was also observed at the mRNA level, and further included high abundance of surface receptors including LAIR1, TLR1, FCRL1, FcεR, interleukin receptors IL4R, IL21R, and transcription factors BTLA, BCL6, BACH2, and FOXO1, and low abundance of surface receptors CD27, CD80, CXCR3, CD70, FAS, CD58, interleukin receptors IL2RA, IL6R, IL10R, IL15RA and BAFF/APRIL receptor TNFRSF13B (Suppl. Table 5). As expected, CD27 mRNA expression was absent in naive B cells and upregulated in MBC subsets. CD27 protein was not detected in any subset despite the CD27-depedent cell-sorting strategy for MBC subsets, indicating the difficulty to detect lower abundant cell-surface proteins in bulk proteomics.

For IgMD MBCs, no proteins were found that were selectively highly expressed, but leukocyte immunoglobulin-like receptor LILRA4 and retinoic acid receptor RXRA transcripts were highly abundant. For IgM, NLRP3 inflammasome transcripts were abundant. For IgA1 and IgA2 no specific highly abundant proteins were found, but IgA2 did express abundant mucosal homing CCR9 transcripts. Also for IgG1, IgG2 and IgG3 no selective abundantly expressed proteins were identified, but IgG2 MBCs differentially expressed, among others, cell adhesion mediator TGFBI, and IgG3 expressed metal transporter SLC11A1 and phospholipase PLCL1. IgG4-specific features are discussed below. Altogether, these data revealed that naive B cell have a unique signature and isotype-defined MBCs share many of the same transcripts and proteins but for several MBC subsets specific upregulated transcripts and proteins could be identified.

### Unique phenotype of IgG4 MBCs

PCA analysis revealed that IgG4 MBCs were most distinct from naive B cells and less similar to other isotype-defined MBCs. Furthermore, K-means clustering analysis identified distinct mRNA and protein expression profiles for IgG4 MBCs: Cluster B (mRNA) and 3 (protein) define mRNA and proteins upregulated in IgG4 MBCs, whereas cluster A (mRNA) and 4 (protein) define downregulated mRNAs and proteins, respectively (Fig. 2D).

Zooming in onto specific mRNAs/protein expression in IgG4 MBCs, a number of selectively down-regulated mRNAs that stand out include FYN and HCK, linked to tyrosine kinase activity (Suppl. Table 5). Indeed, SYK is also found to be selectively downregulated (-0.57), and downregulation SYK was also identified on protein level, confirming the diminished expression of this molecule for IgG4 MBCs relative to all other subsets (Suppl. Table 6). Furthermore, mRNA levels of NF-kB signaling-associated TNFRSF18, CARD11 and EFHD2 are also down-regulated. We also confirmed the previously identified down-regulation of CXCR3 transcripts and CR2 for IgG4 compared to IgG1 MBCs (Fig. S3, Unger et al., 2020). Furthermore, the most downregulated transcript in IgG4 MBCs compared to all other subsets was that of the neonatal Fc receptor (FCGRT/FcRn) important for IgG recycling. Another Fc receptor, FCGR2B, was not amongst the most highly downregulated mRNAs, but still selectively less expressed in IgG4 MBCs (-0.5).

Selective, highly upregulated mRNAs in IgG4 compared to all other subsets (Supp.Table 5) include LDLR, PAX5 repressor TLE4 and chemokine receptor CCR1. Furthermore, SDR16C5 mRNA is highly upregulated (Figure 3B), and this specific upregulation was confirmed at the protein level. SDR16C5, also known as RDH-E2, functions as an oxidoreductase of all-trans-retinol, and serves as a precursor for bioactive metabolites retinal and retinoic acid.

Further transcriptome analysis revealed an upregulation of IL-4R on IgG4 MBCs versus all other IgG subsets and an IgG4-specific upregulation of the IL-13RA1 (Fig. 3D). Cytokines corresponding to these receptors, IL-4 and IL-13, promote class switch towards IgG4 but also to IgE. Surprisingly, IgG4 MBCs display upregulated expression of IGHE transcripts which was not detected on protein level (Fig. 3C-D). Also tolerance-associated cytokine IL-10 may promote class switch towards IgG4. IL-10RA expression levels were similar for IgG4^+^ MBCs compared to other MBC subsets and IL-10 was not detected for IgG4 MBCs on both mRNA and protein level, suggesting that IgG4^+^ MBCs do not produce IL-10 to promote tolerance in an autocrine or paracrine fashion. IL-21R expression was similar for IgG4 MBCs compared to other MBC subsets but lower compared to naive and IgMD^+^ MBCs. IgG4 MBCs also showed higher expression of FcεRII (CD23) compared to other IgG MBC subsets but not compared to naive B cells (CD23, Fig. 3D), suggesting increased capacity for IgG4^+^ MBCs to respond to IgE molecules.

Taken together, novel features for IgG4 MBCs are identified in this study which suggest not only phenotypical differences, but also functional differences for IgG4 MBCs in peripheral blood.

## Discussion

We generated proteomics and transcriptomics datasets of naive and isotype-defined MBC subsets derived from human peripheral blood. Combined proteome and transcriptome analysis revealed that mRNA and protein expression profiles separate B cell subsets according to differentiation status. mRNA and protein expression levels correlated reasonably well for many genes. IgG4 MBCs were identified as being most distinct from naive B cells, which implies that there exist options for selective targeting of this B cell subset, to either enhance immune tolerance or counteract IgG4-switched autoantibody production.

Combined proteome and transcriptome analysis reinforces confidence and allows for robust pathway analysis. These datasets provide a resource for investigation of human naive and isotype-defined MBC identity and function. Due to the low frequency of IgG4 MBCs we could not further define MBCs subsets using additional relevant phenotypic markers such as CD21, CD24, and CD95. In total, we identified 3511 proteins across all subsets, of which 738 proteins were differentially expressed, mostly between MBC subsets compared to naive B cells. Naive and memory B cells exhibit also similar expression profiles for many genes, as both cell types exist in a relatively quiescent state when not actively participating in an immune response. In this study cells were sorted CD38-indicating that they were not activated. These shared genes are linked to a low metabolic rate and the maintenance of a dormant state, enabling both naive and resting memory B cells to survive long-term without the high metabolic demands associated with active antibody production.

Non-switched MBCs are considered more likely to re-enter the GC reaction upon secondary antigen exposure whereas class-switched MBCs are more likely to undergo rapid differentiation toward an antibody-secreting cell type. We observed that non-switched MBCs exhibit a gene expression profile more similar to naive cells suggestive of functional plasticity for this subset and less extensive differentiation compared to class-switched MBCs. Compared to switched MBCs these subsets had higher expression of genes involved in activation, antigen presentation and cytokine signaling, in line with a previous study (3). In that study the non-switched MBC subset was defined as being IgM^+^, comprising heterogenous and undefined IgD expression levels. In the current study we separated non-switched MBC into IgM+IgD^low^ and IgM^+^IgD^+^ MBCs. IgD expression levels are important to include as IgM^+^IgD^+^ MBCs likely reside from the marginal zone compartment and are raised in a different microenvironment shaping different functional properties attributed to differential gene expression.

It is to be expected that different isotype-defined MBC subsets display distinct functional characteristics that influence their ability to expand, migrate and differentiate following activation. Of interest are differences in expression of surface receptors and secreted factors that can modulate responses to antigen, influence their migration capacity and interactions with other cell types in the local environment. IgA2^+^ MBCs expressed higher levels of CCR9, which directs migration to the small intestine (39). CXCR3, important for recruitment to inflammatory sites, was upregulated on all isotype-switched MBCs, but less for IgG4 B cells. The CXCR3 pathway plays a crucial role in recruitment of inflammatory cells in settings of chronic inflammation, for example, in allergic and autoimmune diseases. This suggests less CXCR3-mediated recruitment of IgG4 MBCs into inflamed sited. We did, however, observed upregulation of CCR1 on IgG4 MBCs compared to other MBCs subsets, which may serve a similar function (40,41).

Upon BCR cross-linking the intracellular domain (ITAM) of the Igα/Igβ heterodimer recruits protein tyrosine kinases (PTKs) for initiation and transduction of subsequent signaling events. This signaling cascade can promote different biological outcomes depending on the expressed BCR isotype and additional signals received by the B cell (42–44). ITAM-bound Syk can be phosphorylated thereby lowering the threshold for activation, augmenting B cell survival, proliferation and plasma cell differentiation (45,46). Intriguingly, IgG4 MBCs expressed lower levels of many PTKs, including for example Syk, Lyn, Fyn and HCK, suggesting lowered BCR signaling which may reflect the low frequency of IgG4 MBCs and limited IgG4 plasma cell generation.

Retinol metabolism important in gut IgA MBCs and plays a role in tolerance. Retinol, or vitamin A, is an essential nutrient for a healthy immune system. Retinol serves as a precursor for bioactive metabolites retinal and retinoic acid, the latter is critical for B cell development, proliferation and differentiation. Protein levels of SDR16C5 was upregulated in IgG4 B cells compared to all tested B cell subsets. SDR16C5 functions as an oxidoreductase of all-trans-retinol and serves as a precursor for bioactive metabolites retinal and retinoic acid. If the upregulated expression of SDR16C5 leads to enrichment of retinol-derived bioactive metabolites their function might play are role in the tolerance-associated phenotype of IgG4 MBCs, similar as to what is observed for IgA B cells at mucosae surfaces.

The capacity of MBCs to undergo isotype-switching and selection is essential for understanding secondary immune responses. Previous studies that performed antibody repertoire analysis of peripheral blood B cells revealed that the majority of switch events occur from IgM to IgG1 and IgA1 and switching to IgG2, IgG4, IgE and IgA2 may more often occur through indirect switching (48). IgG4+ MBCs, and to a lesser extent IgG1 MBCs, displayed upregulation of IL-13R, IL-4R and IGHE transcripts but not protein. Note that we performed bulk RNAseq and therefore cannot determine whether the majority of IgG4 MBCs express IGHE transcripts or just a fraction with very high expression levels. It is tempting to speculate that IgG4 MBCs, like IgG1 MBCs (49), may be able to isotype-switch to IgE, which is located down-stream of IgG4 on the IGH locus. IgE antibodies show signs of affinity-maturation, but on average have lower mutation levels as seen for IgG4 (50), making IgG1 and IgM more likely precursor candidates. Alternatively, these may also reflect putative sterile transcripts, observed during *in vitro* cultures of B cells upon addition of IL-4 which promotes switch to IgE, that do not encode for functional IgE antibodies (51–54). We recently observed a profound inhibition of the IL-4R blocking antibody dupilumab on IgG4 switching following repeated mRNA vaccination (57). It is tempting to speculate a prominent role of sequential switching via IgG1 towards IgG4, which was interrupted at the B cell level via IL-4R inhibition. In contrast to IgG4 switching, IgG1 switching *in vitro* is still observed in the absence of IL-4 (23).

In summary, IgG4 MBCs share many features with other MBC subsets, but differential expression of RTKs, STATs, and IL and FcγR surface receptors by IgG4 MBCs indicates additional specialization based on isotype. However, functional studies are necessary to determine the functional consequences. These findings provide insight into IgG4 MBC biology and will enable further studies to interrogate IgG4 MBCs in the context of allergy and IgG4 mediated autoimmune disease. Collectively, combined transcriptomics and proteomics revealed isotype-defined phenotypic specialization of human MBCs and in particular identified a distinct profile for IgG4-switched B cells.

