## Supplemental Figures for "Transcriptome-Proteome analysis of human naive and memory B cell subsets reveal isotype and subclass-specific phenotypes"

### Supplementary materials

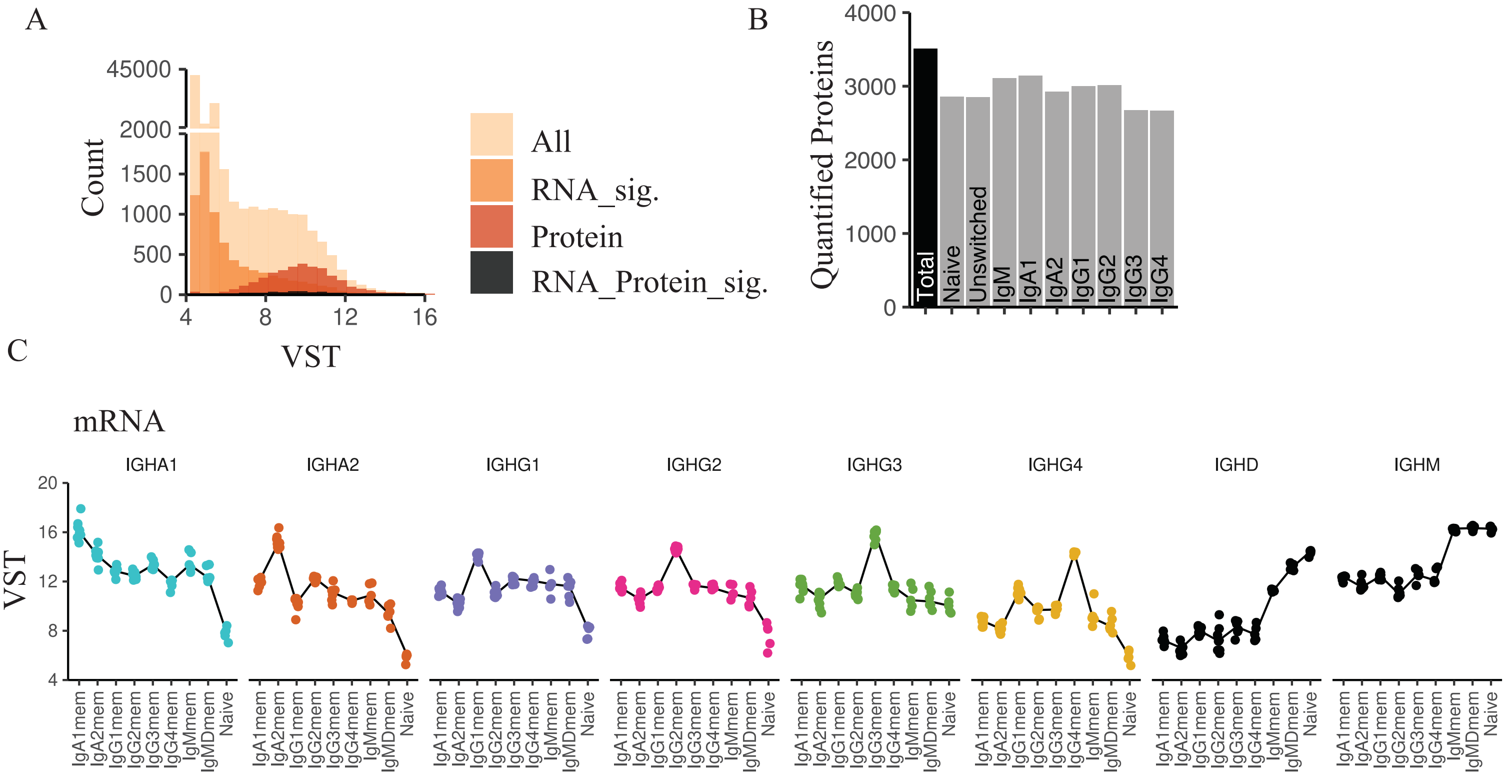


**Figure S1.** RNAseq and protein level comparison. (**A**) Hisogram of RNA expression range depicted as VST. (**B**) Number of quantified proteins for each of the B cell subsets. (**C**) mRNA expression levels for each IGH gene in each B cell subset.

A

| | Down | Up |
|---|---|---|
| NaiveBcells - BmemUnswitched | 28 | 15 |
| NaiveBcells - Bmem_IgM | 85 | 47 |
| NaiveBcells - Bmem_IgA1 | 92 | 54 |
| NaiveBcells - Bmem_IgA2 | 137 | 112 |
| NaiveBcells - Bmem_IgG1 | 102 | 102 |
| NaiveBcells - Bmem_IgG2 | 115 | 76 |
| NaiveBcells - Bmem_IgG3 | 106 | 69 |
| NaiveBcells - Bmem_IgG4 | 152 | 180 |
| BmemUnswitched - Bmem_IgM | 2 | 4 |
| Bmem_IgA1 - Bmem_IgA2 | 1 | 1 |
| Bmem_IgA1 - Bmem_IgG1 | 0 | 3 |
| Bmem_IgA1 - Bmem_IgG2 | 1 | 1 |
| Bmem_IgA1 - Bmem_IgG3 | 0 | 2 |
| Bmem_IgA1 - Bmem_IgG4 | 10 | 54 |
| Bmem_IgA2 - Bmem_IgG1 | 16 | 49 |
| Bmem_IgA2 - Bmem_IgG2 | 0 | 1 |
| Bmem_IgA2 - Bmem_IgG3 | 2 | 2 |
| Bmem_IgA2 - Bmem_IgG4 | 8 | 31 |
| Bmem_IgG1 - Bmem_IgG2 | 2 | 1 |
| Bmem_IgG1 - Bmem_IgG3 | 3 | 4 |
| Bmem_IgG1 - Bmem_IgG4 | 53 | 85 |
| Bmem_IgG2 - Bmem_IgG3 | 0 | 0 |
| Bmem_IgG2 - Bmem_IgG4 | 11 | 34 |
| Bmem_IgG3 - Bmem_IgG4 | 7 | 28 |

B

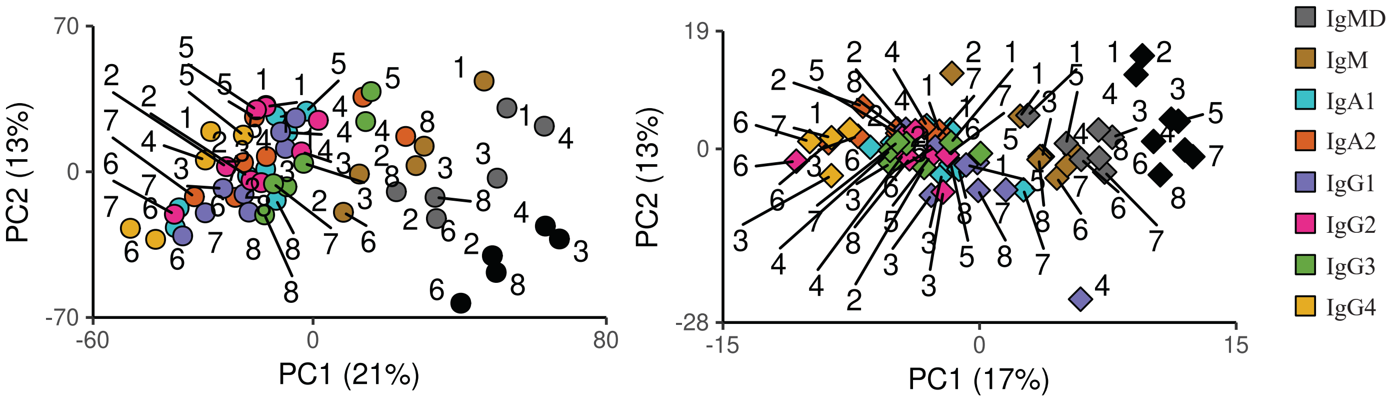


**Figure S2.** (**A**) Pairwise comparison of differentially expressed proteins between all subsets. (**B**) PCA analysis as in Fig. 2A with depiction of donor number per data point.

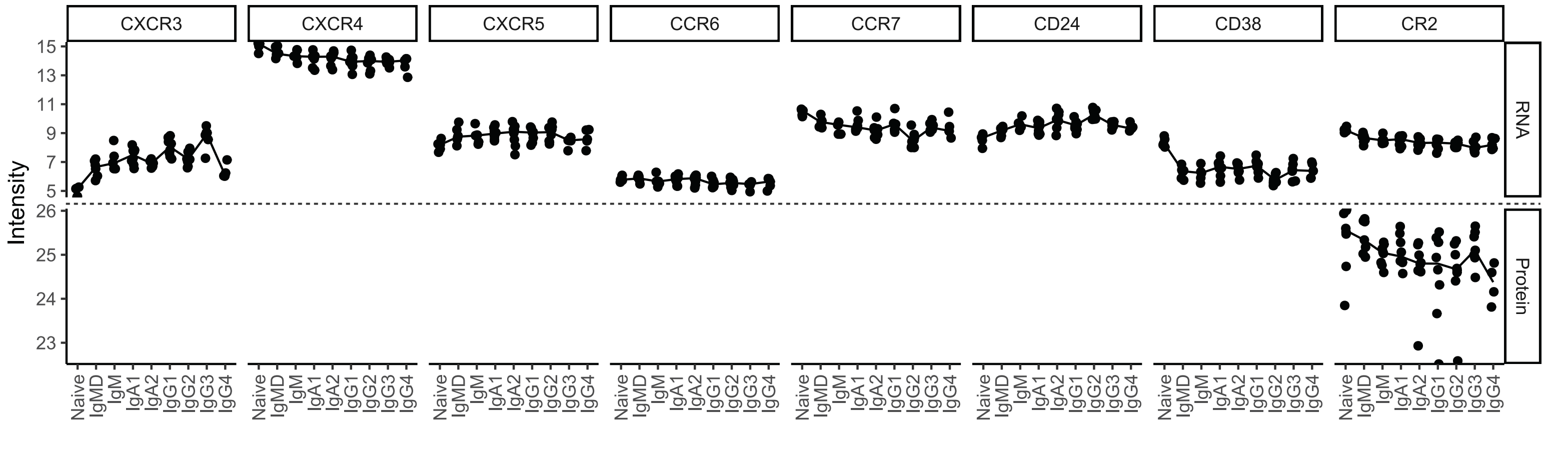


**Figure S3.** mRNA and protein expression levels of previously identified proteins differentially expressed between IgG1 and IgG4 B cells. For many targets no protein was detected.
